# The Role of MAP3K1 in the Development of the Female Reproductive Tract

**DOI:** 10.1101/2023.04.20.537715

**Authors:** Eiki Kimura, Maureen Mongan, Bo Xiao, Jingjing Wang, Vinicius S Carreira, Brad Bolon, Xiang Zhang, Katherine A. Burns, Jacek Biesiada, Mario Medvedovic, Alvaro Puga, Ying Xia

## Abstract

Mitogen-Activated Protein 3 Kinase 1 (MAP3K1) is a dynamic signaling molecule with a plethora of cell-type specific functions, most of which are yet to be understood. Here we describe a role for MAP3K1 in the development of female reproductive tract (FRT). MAP3K1 kinase domain-deficient (*Map3k1^ΔKD^*) females exhibit imperforate vagina, labor failure, and infertility. These defects correspond to a shunted Müllerian duct (MD), the principle precursor of the FRT, in embryos, while they manifest as a contorted caudal vagina with abrogated vaginal-urogenital sinus fusion in neonates. In epithelial cells, MAP3K1 acts through JNK and ERK to activate WNT, yet *in vivo* MAP3K1 is crucial for WNT activity in mesenchyme associated with the caudal MD. Expression of *Wnt7b* is high in wild type, but low in *Map3k1* knockout MD epithelium and MAP3K1-deficient keratinocytes. Correspondingly, conditioned media derived from MAP3K1-competent epithelial cells activate TCF/Lef-luciferase reporter in fibroblasts, suggesting that MAP3K1-induced factors released from epithelial cells trans-activate WNT signaling in fibroblasts. Our results reveal a temporal-spatial and paracrine MAP3K1-WNT crosstalk contributing to MD caudal elongation and FRT development.

**Highlights:**

- MAP3K1 deficient female mice exhibit imperforate vagina and infertility
- Loss of MAP3K1 kinase activity impedes Müllerian duct (MD) caudal elongation and fusion with urogenital sinus (UGS) in embryogenesis
- The MAP3K1-MAPK pathway up-regulates WNT signaling in epithelial cells
- MAP3K1 deficiency down-regulates Wnt7b expression in the MD epithelium and prevents WNT activity in mesenchyme of the caudal MD

## Introduction

The development of the mammalian reproductive system is a complex process with distinct sex-independent and -dependent phases. Prior to sexual differentiation, the reproductive tract precursors, known as the Müllerian (or paramesonephric) ducts (MDs) and the Wolffian (or mesonephric) ducts (WDs), develop in both male and female embryos. These ducts originate from the intermediate mesonephros and elongate caudally towards the urogenital sinus (UGS) (Orvis and Behringer, 2007). In mice, the WD reaches and fuses with the UGS at embryonic day (E) 11.5 before developing into the internal genitalia of males, whereas the MD fuses with the UGS later at E13.5 and ultimately contributes to the internal genitalia of females (Mullen and Behringer, 2014). The WD also has a role in guiding the development of the MD (Orvis and Behringer, 2007).

Sexual differentiation in mice occurs at E13.5. At this stage, gonads in male embryos begin to produce anti-Müllerian hormones and androgens. These hormones are responsible for MD regression but WD stability. The stabilized WDs subsequently differentiate into epididymides, vas deferens and seminal vesicles of the male reproductive tract. In female embryos, the absence of these hormones results in regression of the WDs. Their MDs, on the other hand, continue to develop into the FRT derivatives, i.e. the oviduct, uterine horns, cervix, and the upper vagina (Roly et al., 2018). Prior to puberty, the vagina is composed of a solid epithelial cord derived from MD and UGS epithelia. The vagina becomes canalized in adulthood, resulting in formation of the vagina orifice at approximately 24-28 days depending on the mouse strain (Kurita, 2010).

The molecular mechanisms underlying MD formation and development remain poorly understood, although studies in mice have identified a number of genes that play pivotal roles (Orvis and Behringer, 2007; Parr and McMahon, 1998; Prunskaite-Hyyrylainen et al., 2016). Some of the genes identified in mice, such as *Wnt4* and *Hoxa10*, are found mutated in FRT anomaly cases in humans, underscoring conserved genetic control of FRT development across the species (Biason-Lauber et al., 2004; Cheng et al., 2011). To date, gene mutations are found in a small number of FRT anomalies, whereas the genetic etiology for the majority of FRT anomaly cases are yet to be identified (Laronda et al., 2013).

The mitogen-activated protein kinase kinase kinase 1 (MAP3K1) is a member of the MAP3K superfamily, consisting of 19 protein kinases (Uhlik et al., 2004). The primary role of the MAP3Ks is the regulation of MAPK activation, which in turn participates in diverse cellular activities, such as gene expression and the cell proliferation, migration, survival and death (Craig et al., 2008). MAP3K1 is a large protein consisting of 1500 amino acids and has several well-defined functional domains, such as the Kinase Domain (KD), Plant Homeodomain (PHD), Guanine Exchange Factor (GEF) (Wang et al., 2021a). The different domains confer divergent functions to this protein. For example, the PHD domain possesses E3 ubiquitin ligase activity that mediates protein degradation, whereas the KD is responsible for stimuli-specific activation of the MAP2K-MAPK cascades (Lu et al., 2002; Xia et al., 1998).

Analyses of *Map3k1* mutant mice have identified a plethora of functions for MAP3K1 in embryo survival, immune cell maturation, and the development of sensory organs, such as eye and ear (Charlaftis et al., 2014; Gallagher et al., 2007; Labuda et al., 2006; Yousaf et al., 2015; Yujiri et al., 2000; Zhang et al., 2003). In addition, human genomic studies have uncovered a strong association of *MAP3K1* mutations with sex development and differentiation (Das et al., 2013; Granados et al., 2017; Pearlman et al., 2010). Collective genomics data worldwide have identified germline gain-of-function *MAP3K1* variants in 13-18% of individuals with 46, XY disorder of sex development (DSD). The 46, XY DSD is a disease defined as abnormal sexual differentiation where patients with a male genotype is characterized by a female phenotype with respect to sexual features (Allen, 2009; Ostrer, 2014).

In light of these findings in humans, the roles of MAP3K1 in male sexual development have been explored using *Map3k1^ΔKD^* mice (Warr et al., 2011). The *Map3k1^ΔKD^* genotype is a knockin/knockout strain in which the exon coding for the MAP3K1 kinase domain is replaced by a β-galactosidase (β-GAL) gene (Xia et al., 2000). The resultant *Map3k1^ΔKD^* allele expresses a kinase domain deficient MAP3K1-β-GAL fusion protein. Through the examination of ß-GAL activity in the *Map3k1^+/ΔKD^* embryos, abundant MAP3K1 expression was detected in the gonads, testis, reproductive tracts and mesonephric tubules in the E11-E13.5 embryos without sexual dimorphism. The expression pattern was consistent with the anticipated role of MAP3K1 in sex organ development; however, the *Map3k1* homozygous knockout embryos had no major defects in testis development and the adult males were fertile. These results argue that MAP3K1 kinase activity is dispensable for male sexual development (Warr et al., 2011).

The nature of the mouse and the human *MAP3K1* mutations is fundamentally different. In *Map3k1^ΔKD^* mice, an 800 bp deletion that removes the KD coding sequences results in a loss-of-function mutation. In contrast, human 46, XY DSD patients bear one or several nucleotide variants that spread throughout the *MAP3K1* gene. The most common non-synonymous variants are associated with gain-of-function mutations involving the GEF domain coding region (Pearlman et al., 2010). The GEF mutants display enhanced MAP3K1 signaling activity, including stronger binding with co-factors (i.e. RHOA and AXIN), increased activation of the downstream targets (i.e. MAPKs), and activation of the WNT-β-catenin pathway (Chamberlin et al., 2019; Loke and Ostrer, 2012; Loke et al., 2014; Pearlman et al., 2010; Xue et al., 2019). Taken together, these data indicate that the gain-of-function *MAP3K1* variants impede male but potentiate female sexual features.

In this study, we examined *Map3k1^ΔKD^* mice and identified a role for MAP3K1 in the development of the female reproductive tract. Our data show that MAP3K1-WNT paracrine crosstalk is a signaling mechanism that contributes to FRT development and that *Map3k1* loss-of-function mutation is a likely genetic etiology for FRT anomalies underlying female infertility.

## Results

### Missing MAP3K1 KD impairs FRT formation and function

We examined the fertility of adult C57/BL6 mice with wild type, *Map3k1^+/ΔKD^* and *Map3k1^ΔKD/ΔKD^*genotypes. The male *Map3k1^ΔKD/ΔKD^* mice displayed normal sexual organ development and fertility, similar to that reported previously (Warr et al., 2011). In contrast, a significant proportion of *Map3k1^ΔKD/ΔKD^*females were infertile due to a lack of vaginal opening, a phenomenon known as an imperforate vagina (IPV) (Fig. 1A). Out of 38 *Map3k1^ΔKD/ΔKD^* females examined, 45% displayed IPV and over time developed lower abdominal distention. The swollen abdomen of the *Map3k1^ΔKD/ΔKD^* IPV mice resulted from bilateral enlargement of the uterine horns due to retention of protein-rich fluid, exfoliated keratin, leukocytes, and a few necrotic epithelial cells (Figs. 1B and 1C). Medical designation for this condition is hydrometrocolpos. The wild type uteri displayed normal histology, two layers of myometrium, endometrium containing stromal and epithelial cells and numerous epithelial lined glands. In contrast, the uteri of *Map3k1^ΔKD/ΔKD^* IPV mice displayed thin myometrial layers, minimal stromal cells in the endometrium, and lacked visible glands. Epithelial cells were only seen in the uterine lumen and not in glands. Histological examination of the *Map3k1^ΔKD/ΔKD^*IPV mice confirmed that the caudal end of the vaginal epithelium was not connected to the surface of the perineal skin, thereby preventing vaginal canalization at puberty (Fig. 1D). Without a vaginal opening (perforate), the IPV mice could not be impregnated.

**Figure 1.**
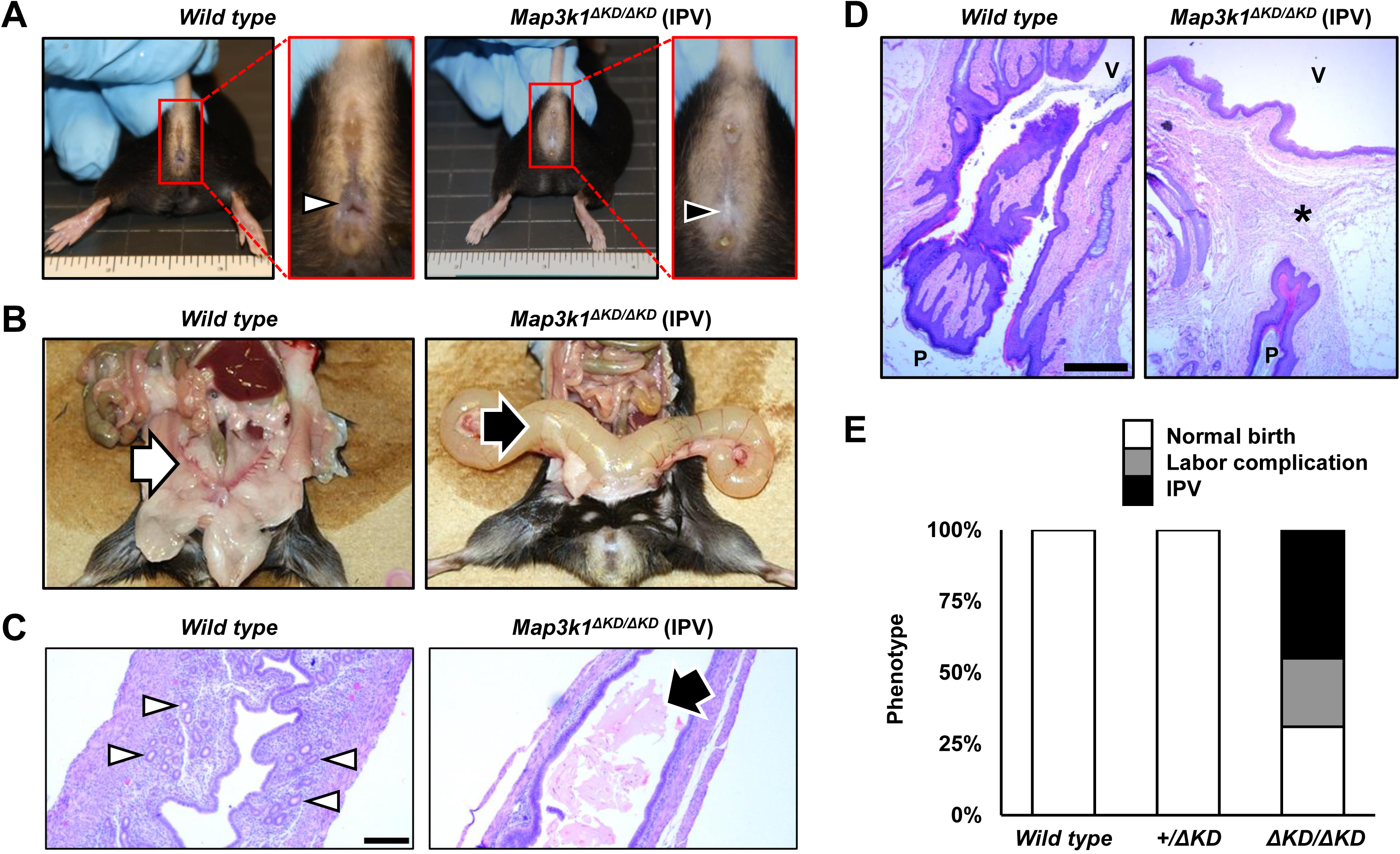
*Map3k1_ΔKD/ΔKD_* female mice have an imperforate vagina (IPV) phenotype and reduced infertility. Multiple 1-to 3-month-old wild type and *Map3k1^ΔKD/ΔKD^* females were examined for reproductive defects. (A) Arrowheads point to the patent vaginal opening in wild type and its absence in the IPV *Map3k1^ΔKD/ΔKD^* females, (B) Arrows point to narrow uterine horns in wild type compared to dilated uterine horns in *Map3k1^ΔKD/ΔKD^* females with IPV. Histology with H&E staining showed (C) a narrow uterine lumen with thick wall and prominent endometrial glands (arrowheads) in wild type compared to the dilated lumen containing proteinaceous secretory material admixed with exfoliated cellular debris (arrow), lined by a thinner uterine wall with fewer endometrial glands in *Map3k1^ΔKD/ΔKD^*females; and (D) a canalized caudal vagina connected with the perineal skin in the wild type in contrast to IPV with vaginal epithelium separated from the perineal skin by mesenchymal tissues (*) in the *Map3k1^ΔKD/ΔKD^* females with IPV. P, surface of perineal skin; V, vagina. (E) Quantification of female reproductive activity showed that while all of wild type (n=70) and *Map3k1^+/ΔKD^*(n=97) had normal pregnancy and fertility, 77% of the *Map3k1^ΔKD/ΔKD^*mice (n=38) had abnormal reproduction due to either IPV or labor complications yielding no live pups. Scale bars, 200 and 500 µm in C and D, respectively.

Some *Map3k1^ΔKD/ΔKD^* females did not have the IPV phenotype. They could become pregnant, but half of them failed to deliver live offspring (Fig. 1E). The other half were fertile with no apparent reproductive defects, indicating a properly functioning hypothalamic gonadial axis. Neither the wild type (n=70) nor the *Map3k1^+/ΔKD^*(n=97) mice had FRT defects or labor complications. Thus, the *Map3k1^ΔKD^* allele is autosomal recessive in reducing reproductive health and causing infertility in female mice.

### The developmental origins of the FRT defects

To determine whether the FRT abnormalities observed in *Map3k1^ΔKD/ΔKD^* adults were congenital, we examined neonates. Using anti-keratin (K) 14 for basal epithelial cells and whole organ staining, we detected the expected symmetrical V-shaped vagina structure and vagina-UGS fusion zone in the *Map3k1^+/ΔKD^* females (Fig. 2A; Suppl video 1-3). In contrast, the vagina was largely distorted or absent due to truncated MDs failing to reach the UGS in the *Map3k1^ΔKD/ΔKD^*female. Immunohistochemistry using anti-β-catenin that labels epithelial cells, we found that compared to the *Map3k1^+/ΔKD^*, the *Map3k1^ΔKD/ΔKD^* pups displayed a distorted or missing vagina in the caudal FRT sections (Fig. 2B). Importantly, 8 out of 9 (89%) *Map3k1^ΔKD/ΔKD^* pups had distorted vagina structure (Fig. 2C), a ratio higher than that of infertile adults (approximately 70%), indicating that some, presumably modest, congenital defects can recover during postnatal development.

**Figure 2.**
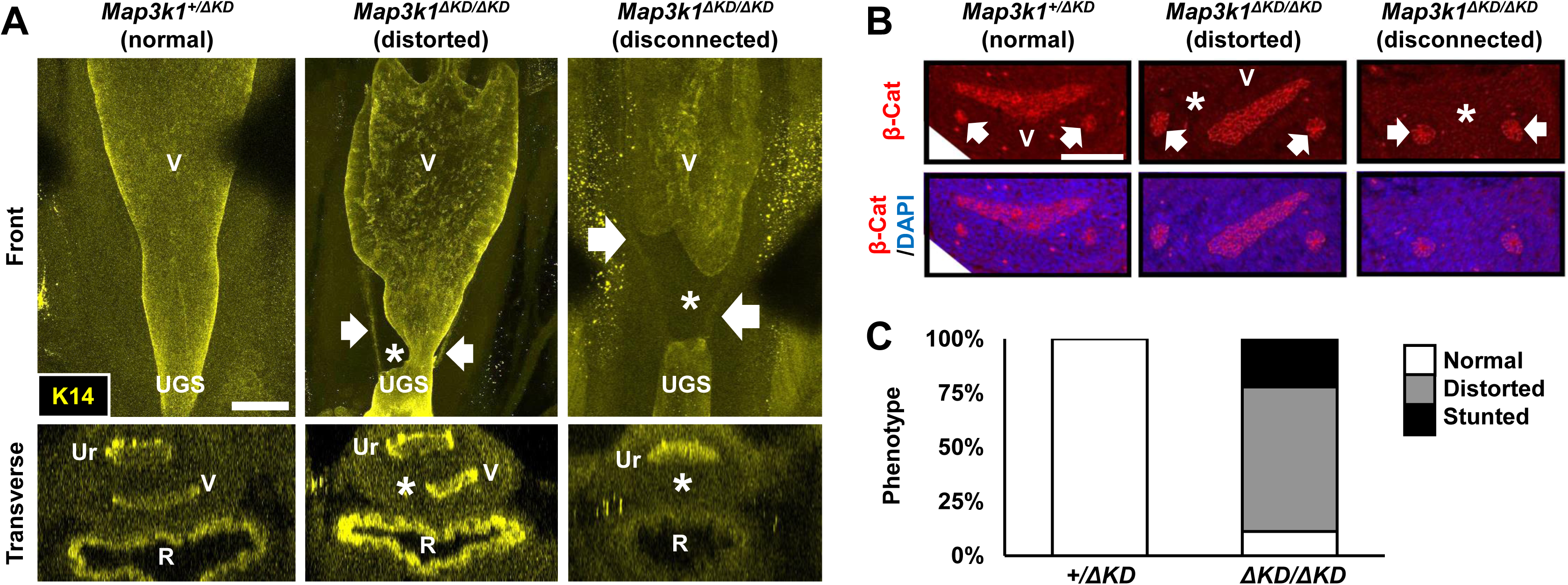
Vagina-urogenital sinus (UGS) fusion defects in the *Map3k1_ΔKD/ΔKD_* neonates. (A) The caudal end of the vagina and vagina-UGS fusion site of *Map3k1^+/ΔKD^* and *Map3k1^ΔKD/ΔKD^* neonates were examined by whole-mount immunostaining with anti-Keratin 14 (K14) that labels the epithelium lining the reproductive tract, and 3D images were captured with a confocal microscope. Representative longitudinal and transverse images are shown. (B) Transverse sections of the caudal reproductive tract of neonates were assessed by immunohistochemistry with anti-β-Catenin (β-Cat) to label epithelial cells. Two representative defect phenotypes—distorted and stunted vaginas—observed in the *Map3k1^ΔKD/ΔKD^*mice are shown. (C) The incidences of the normal, distorted and stunted vagina found in 3 *Map3k1^+/ΔKD^* and 9 *Map3k1^ΔKD/ΔKD^*neonates. Arrows, Wolffian ducts; R, rectum; UGS, urogenital sinus; Ur, urethra; V, vagina; asterisks mark the stunted and distorted vagina in the *Map3k1^ΔKD/ΔKD^* pups. Scale bars, 200 and 100 µm in A and B, respectively.

To further trace the developmental origin of the FRT defects, we performed whole mount immunostaining of E15.5 embryos using anti-K8, a simple epithelia marker. The 3-dimensional (3D) images showed that the WDs and the MDs extended and fused with the UGS in the expected manner in both female and male *Map3k1^+/ΔKD^* embryos (Roly et al., 2018) (Fig. 3A; Suppl videos 4-7). In the *Map3k1^ΔKD/ΔKD^*embryos, the WDs also extended and fused with the UGS, while the MDs were visibly truncated and MD-UGS fusion was abnormal or lacking. The caudal MD abnormality was observed in both female and male embryos.

**Figure 3.**
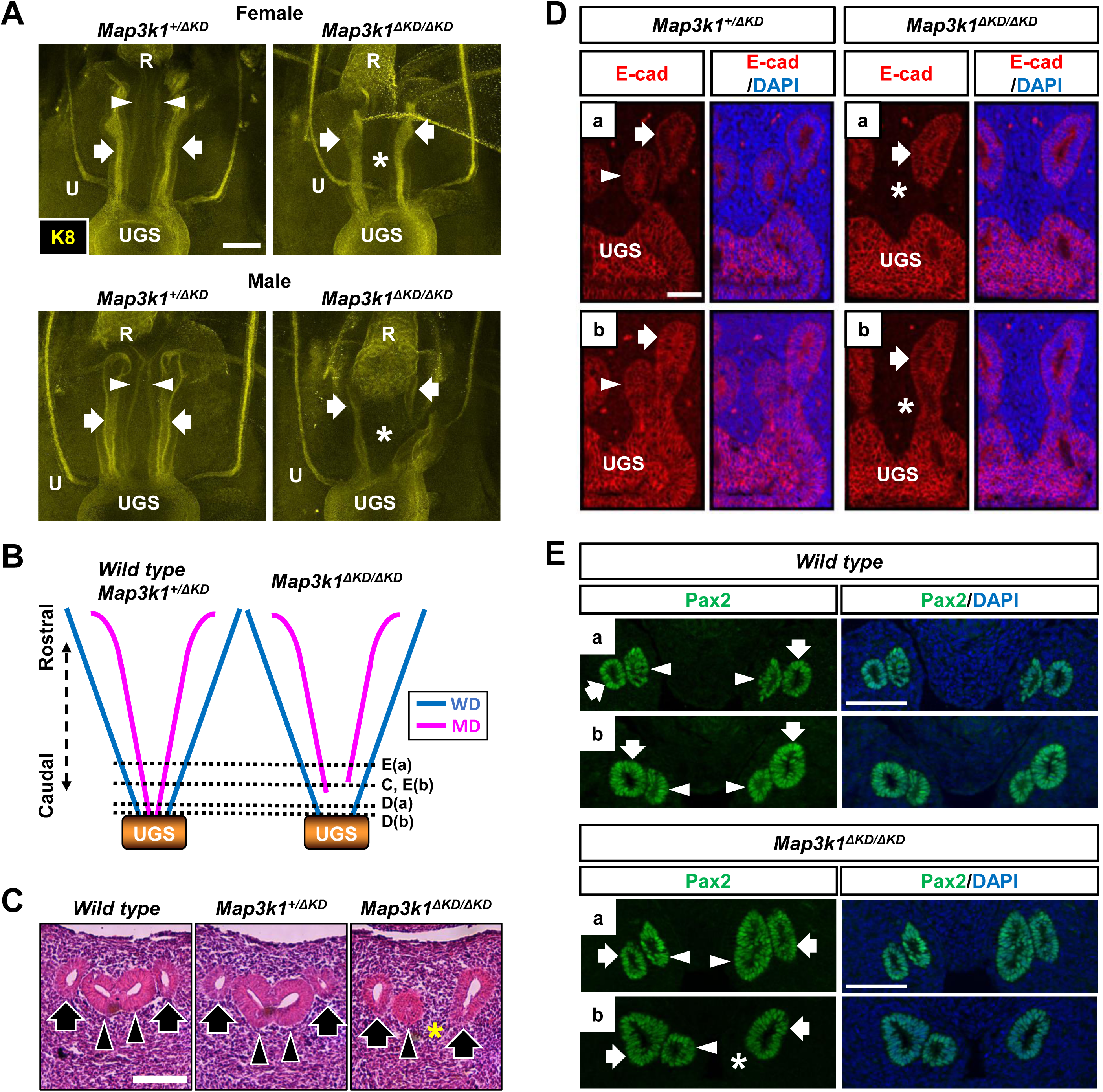
MAP3K1 is required for Müllerian duct (MD) caudal elongation and establishment of MD-urogenital sinus (UGS) fusion. Reproductive tracts of female and male *Map3k1^+/ΔKD^* and *Map3k1^ΔKD/ΔKD^* E15.5 embryos were processed for whole mount staining with anti-Keratin 8 (K8) that labels the simple epithelia, and 3D images were captured with a confocal microscope. (A) Representative longitudinal images of the developing urogenital system. (B) A diagrammatic illustration of the observed rostral and caudal reproductive tracts, where dotted lines mark the relative positions of the serial transverse sections shown in panels C, D and E. The transverse sections at E15.5 are processed for (C) H&E staining and (D) immunohistochemistry with anti-E-cadherin (E-cad), a marker located on the epithelial cell membrane, (E) Transverse sections of the rostral and caudal reproductive tracts in wild type and *Map3k1^ΔKD/ΔKD^* embryos at E13.5 were processed by immunohistochemistry to detect anti-Pax2, a marker of developing reproductive tract epithelium. WD, Wolffian duct (arrows, and blue lines in B); MD, Müllerian duct (arrowheads, and red lines in B); U, ureter; R, rectum. Asterisks mark the absence of MD in *Map3k1^ΔKD/ΔKD^* embryos. Scale bars, 100 µm in A and E, and 50 µm in C and D.

To validate the caudal MD defects in the *Map3k1^ΔKD/ΔKD^*embryos, we performed a histological evaluation of transverse serial sections of the reproductive tract (Fig. 3B). Using H&E staining, we detected the expected two WDs and two MDs at the caudal sections of the reproductive tracts in wild type and *Map3k1^+/ΔKD^* embryos (Fig. 3C). In contrast, we found two WDs but one MD in the corresponding sections of the *Map3k1^ΔKD/ΔKD^*embryos. Immunohistochemistry for E-cadherin that labels epithelial cells confirmed these observations. While the caudal end of the MD was fused with the UGS in *Map3k1^+/ΔKD^* embryos, it was disconnected from UGS in the *Map3k1^ΔKD/ΔKD^* embryos (Fig. 3D). The MD defects were observed in all of the 14 *Map3k1^ΔKD/ΔKD^* embryos but in none of the 5 wild type and 19 *Map3k1^+/ΔKD^* embryos.

The defects observed in the E15.5 *Map3k1^ΔKD/ΔKD^* embryos could be due to insufficient proliferation or excessive apoptosis that led to breaking up of the already established MD-UGS connections. Cell proliferation (by EdU labeling) and apoptosis (by TUNEL assay) were both detected in the reproductive tracts of animals regardless of genotype, and their levels were not quantitatively different in *Map3k1^+/ΔKD^* and *Map3k1^ΔKD/ΔKD^*embryos (Suppl Fig. 1A-D). These findings indicate that aberrant proliferation and apoptosis are not responsible for the MD defects in the knockout mice.

To evaluate whether the defects were due to a failure of MD caudal elongation, we examined embryos of E13.5, a developmental stage when the MDs are just beginning to establish contact with the UGS (Mullen and Behringer, 2014). Examination of serial transverse sections of the embryonic reproductive tract epithelium (Pax2-positive) revealed that the rostral MDs were identical in embryos of different genotypes, but the caudal MDs were strikingly different between wild type and *Map3k1^ΔKD/ΔKD^* embryos. While the caudal MDs in the wild type animals were fully extended to reach the UGS, either one or both of MDs were stunted in the *Map3k1^ΔKD/ΔKD^* embryos (Fig. 3E). Our data suggest that MAP3K1 kinase activity is required for caudal MD elongation to successfully establish the MD-UGS connection.

### MAP3K1 regulates MAPK activity in the embryonic reproductive tract

Because the *Map3k1^ΔKD^* allele encodes a MAP3K1-β-GAL fusion protein that is controlled under the endogenous *Map3k1* promoter, we performed whole mount X-gal staining of E13.5 embryos to trace MAP3K1 expression. Compared to the lack of X-gal in the wild type embryos, abundant X-gal positive cells (blue) were detected in the reproductive tract precursors of *Map3k1^+/ΔKD^* embryos (Fig. 4A). The X-gal intensity was particularly strong on the luminal side of the WD and MD epithelium, though sporadic and weak X-gal staining was also detectable in the surrounding mesenchyme. These data indicate that MAP3K1 is concentrated in epithelial tissue of the nascent reproductive tract.

**Figure 4.**
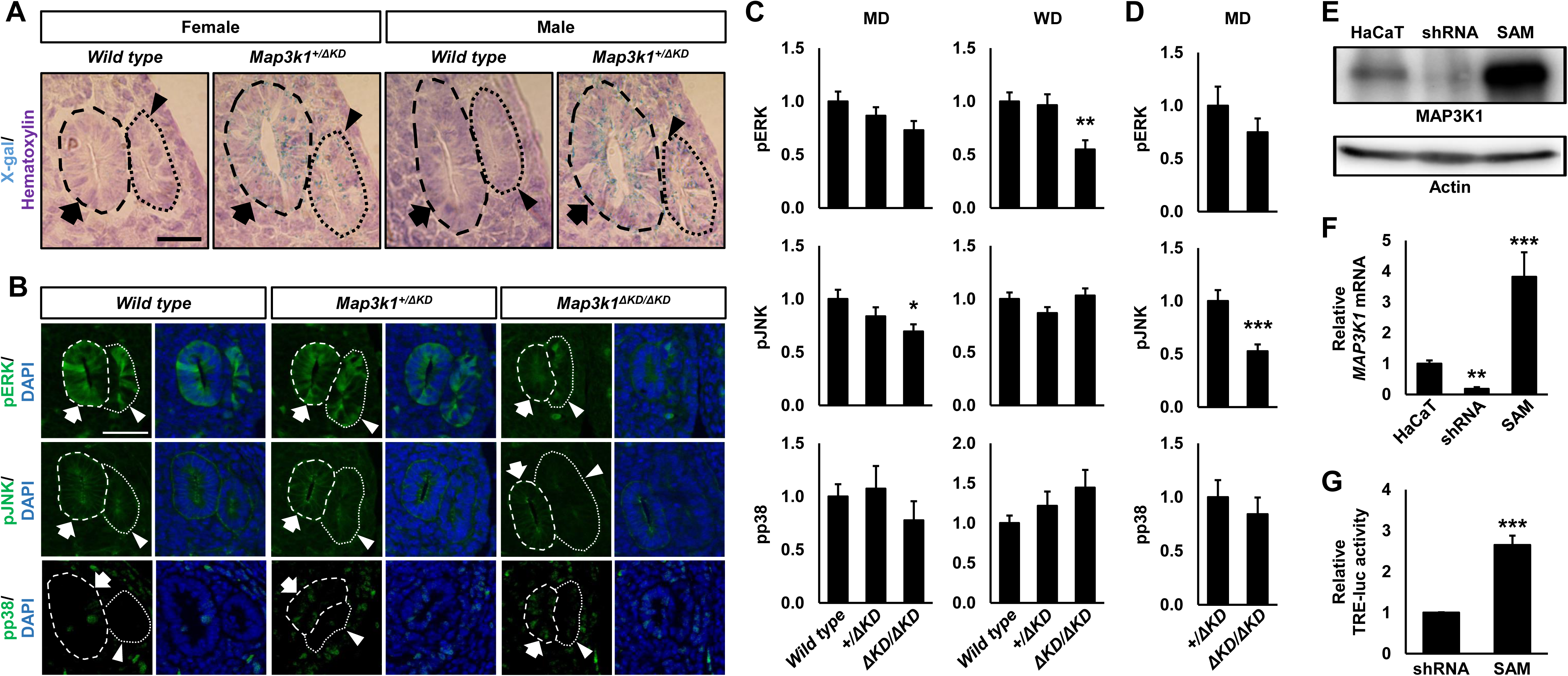
MAP3K1 expression and signaling activity in the reproductive tract. (A) Wild type and *Map3k1^+/ΔKD^* embryos at E13.5 were processed for whole mount X-gal staining; representative sections were photographed with a bright-field light microscope. (B) Tissue sections of the reproductive tract from wild type, *Map3k1^+/ΔKD^* and *Map3k1 ^ΔKD/ΔKD^* embryos were processed to immunolabel pJNK, pERK or pp38; representative images were captured using a fluorescent microscopy. Intensity of the fluorescent signal in the Müllerian ducts (MD) and Wolffian ducts (WD) were quantified at (C) E13.5 and (D) E15.5. At least 3 tissue sections/embryo and 3 embryos/genotype were examined at each time point. The WD (arrows) and MD (arrowheads) were marked with dotted circles. Scale bars, 50 μm in A and B. The HaCaT cells and genetically modified derivative cell lines (shRNA, for MAP3K1 knockdown, and SAM, for MAP3K1 overexpression) were validated by (E) Western blotting using anti-MAP3K1 and (F) qRT-PCR. Compared to the levels in HaCaT cells, MAP3K1 expression was significantly decreased in shRNA cells but significantly increased in SAM cells. (G) TRE-luc activities were determined after adjusting for the transfection efficiency with β-GAL. Statistical analyses of triplicate samples show significantly different TRE-luc activities. Values are mean ± SEM. \**p*<0.05, \*\**p*<0.01 and \*\*\**p*<0.001 are considered significantly different compared to values in wild type or *Map3k1^+/ΔKD^*, or to that in control cells (HaCaT or shRNA).

To assess whether MAP3K1 KD deficiency affects MAPKs, we examined MAPK phosphorylation in wild type, *Map3k1^+/ΔKD^* and *Map3k1^ΔKD/ΔKD^* embryos. The phosphorylation of Extracellular-Regulated Kinase (ERK) and Jun N-terminal Kinase (JNK) was detected in WD and MD epithelium, with the signal intensity particularly strong on the luminal side of the ducts (Fig. 4B). The phosphorylation of p38 MAPK was relatively weak and virtually undetectable in the E13.5 embryos of all genotypes. Compared to that in wild type and *Map3k1^+/ΔKD^* embryos, pERK in *Map3k1^ΔKD/ΔKD^* embryos was significantly reduced in WDs (Fig. 4C). pJNK, on the other hand, was found to be decreased in the MDs, but not the WDs, of the *Map3k1^ΔKD/ΔKD^* embryos. Similar observations were made in the E15.5 embryos: the MDs in the *Map3k1^ΔKD/ΔKD^* embryos displayed a significant reduction of pJNK (Fig. 4D).

To validate the activity of MAP3K1-JNK/ERK signaling axis in epithelial cells, we examined the activation of the AP-1 transcription factors, which bind to the TPA-responsive element (TRE) to activate gene expression (Karin et al., 1997). The TRE-luciferase (TRE-luc) reporter was transfected into human epithelial HaCaT cells genetically-modified to have increased MAP3K1 expression by CRISPR/Cas9 Synergistic Activation Mediator system (SAM) and decreased expression by small hairpin RNA knockdown (shRNA) (Fig. 4E). Compared to the parental HaCaT, the SAM cells had a nearly 8-fold increase while the shRNA cells had an 80% reduction of MAP3K1 expression (Fig. 4F). The TRE-luc, accordingly, exhibited a nearly 3-fold increase in SAM versus shRNA cells (Fig. 4G).

### MAP3K1 crosstalks with the canonical WNT pathways

The *in vivo* and *in vitro* findings suggest that MAP3K1 regulates the JNK/ERK pathway, but how these pathways affect FRT development remains unknown. MAP3K1 and MAPKs are known to crosstalk with the canonical WNT pathway that is implicated in many aspects of sexual organ development (Haraguchi et al., 2017; Luo et al., 2003; Prunskaite-Hyyrylainen et al., 2016; Zhang et al., 2021; Zhang et al., 1999). We therefore examined WNT signaling in MAP3K1-competent and -deficient cells. We performed RNA-seq using SAM and shRNA HaCaT cells, as well as keratinocytes derived from wild type and *Map3k1^ΔKD/ΔKD^* mice (Wang et al., 2021b). A total of 3544 genes were differentially expressed by >2-fold between shRNA and SAM cells, and 2082 genes differentially expression in wild type and *Map3k1^ΔKD/ΔKD^* mouse keratinocytes. Ingenuity pathway analyses of the differentially expressed genes demonstrated that MAP3K1-deficiency leads to down regulation of several WNT-related pathways (Fig. 5A). To validate the MAP3K1-WNT relationships, we transfected the WNT pathway reporter, TCF/Lef-luc, into SAM and shRNA cells. Compared to that in SAM cells, luciferase activities in shRNA cells were reduced by 50% (Fig. 5B). Moreover, several well-known WNT target genes, including *TCF4*, *FZD7, FOSL1 and JUN,* were expressed at lower levels in shRNA cells than in SAM cells (Fig. 5C).

**Figure 5.**
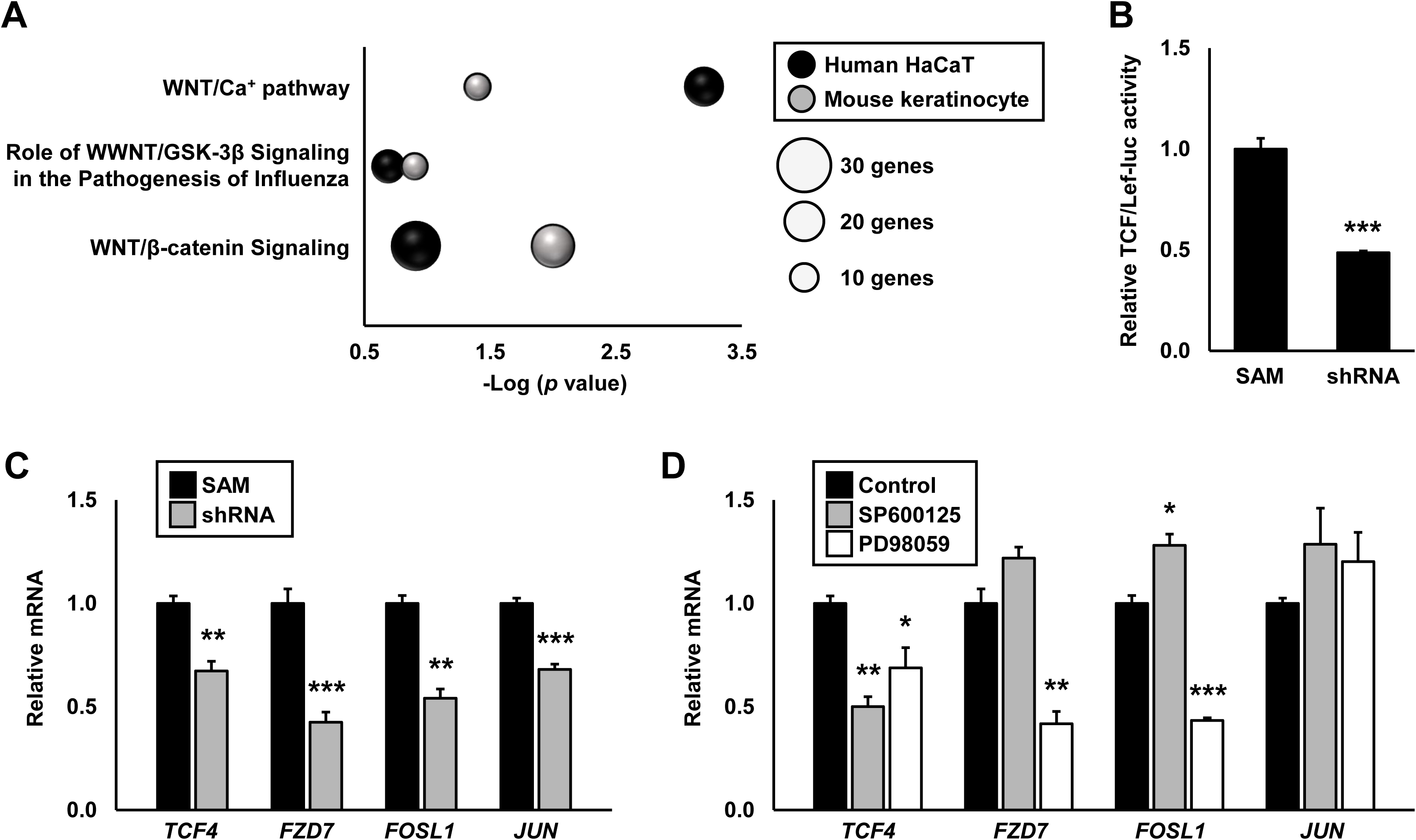
MAP3K1 regulates WNT signaling through MAPK activity. (A) Ingenuity Pathway Analyses (IPA) of differentially expressed genes in SAM-HaCaT and shRNA-HaCaT cells and wild type and *Map3k1^ΔKD/ΔKD^*mouse keratinocytes revealed that MAP3K1 deficiency down-regulated WNT signaling. (B) Differential luciferase activities were detected in SAM and shRNA cells transiently transfected with TCF/Lef-luc (a WNT reporter construct). Expression of candidate WNT target genes was examined by qRT-PCR in (C) SAM and shRNA cells and in (D) SAM cells in the absence and presence of JNK (SP600125) and ERK (PD98059) inhibitors. Results represent data from at least 3 replicate experiments. Values are mean ± SEM. \**p*<0.05, \*\**p*<0.01 and \*\*\**p*<0.001 are significantly different between SAM and shRNA cells or control and inhibitor-treated samples.

To assess whether the effects of MAP3K1 on WNT signaling were mediated through the MAPKs, we treated the SAM cells with MAPK inhibitors and examined WNT target gene expression. Inhibition of JNK by SP600125 did not change the expression of *FZD7* and *JUN*, slightly increased the expression of *FOSL*, but significantly reduced the expression of *TCF4* (Fig. 5D). Inhibition of ERK by PD98059 had no effects on the expression of *JUN*, and down-regulated the expression of *TCF4*, *FOSL* and *FZD7*. These observations suggest that MAP3K1-induced activation of the WNT pathway is mediated partially through JNK and ERK.

### MAP3K1 activates WNT signaling in a paracrine fashion

To examine the MAP3K1-WNT crosstalk *in vivo*, we introduced TCF/Lef:H2B-GFP in wild type, *Map3k1^+/ΔKD^* and *Map3k1^ΔKD/ΔKD^* mice. The TCF/Lef:H2B-GFP links TCF/Lef, a WNT-inducible enhancer, with the gene for green fluorescent protein (GFP), such that WNT activation leads to GFP expression (Ferrer-Vaquer et al., 2010). Examination of E15.5 embryos found temporal-spatial distinct WNT activities along the reproductive tracts. The rostral reproductive tracts had evident GFP signal in the E-cadherin (epithelium) and Pax2 (developing reproductive tract epithelium) double-positive WD and MD epithelium but little if any GFP positivity in the surrounding mesenchyme (Figs. 6A and 6B). The intensity of GFP was not different among the *Map3k1* genotypes (Figs. 6D and S2A). In contrast, the caudal reproductive tract exhibited a distinctive genotype-specific WNT activity. While all embryos regardless of the genotypes had similar GFP in the WD and MD epithelium (Fig. S2B), only the wild type and the *Map3k1^+/ΔKD^* embryos, but not the *Map3k1^ΔKD/ΔKD^* embryos, had GFP positivity in the surrounding mesenchyme (Figs. 6A, 6C and 6D). Similar observations were made in E13.5 embryos, where in the caudal reproductive tract WNT activity was reduced in mesenchyme, but not in the epithelium, of *Map3k1^ΔKD/ΔKD^* compared to wild type embryos (Figs. S2C – S2F).

**Figure 6.**
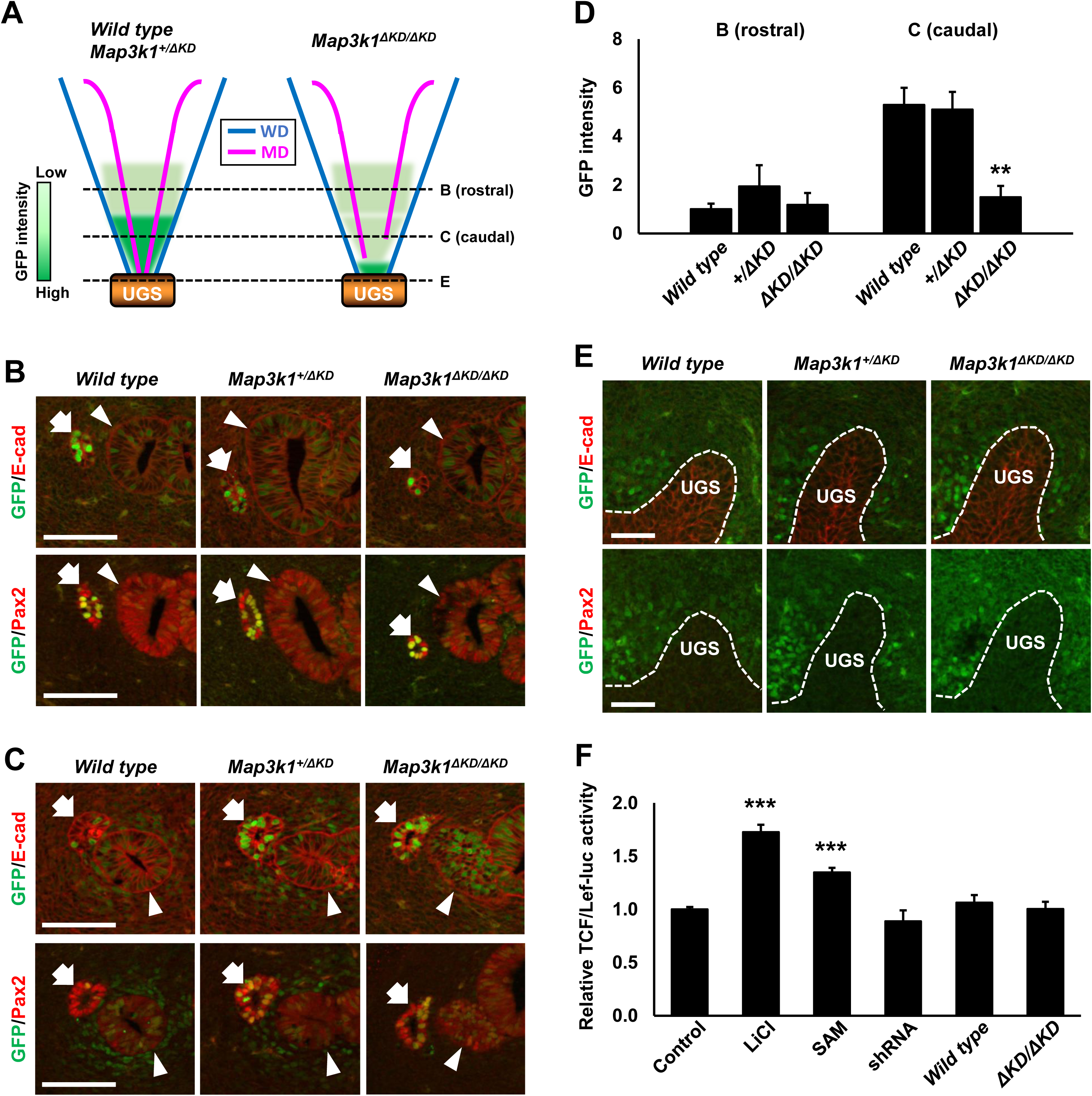
MAP3K1 deficiency abolishes WNT signaling in the mesenchyme associated with the caudal reproductive tract. Inclusion of the WNT reporter (TCF/Lef:H2B-GFP) in wild type, *Map3k1^+/ΔKD^* and *Map3k1^ΔKD/ΔKD^*E15.5 embryos were processed to immunolabel Pax2 and E-cadherin. (A) Diagrammatic illustration of the observed GFP signal intensity in the rostral and caudal reproductive tract and the urogenital sinus (UGS). Dotted lines mark the relative positions of the transverse sections shown in panels B, C and E. Representative images of the (B) rostral and (C) caudal reproductive tracts and the (E) UGS were captured with a fluorescent microscope. Scale bars,100 μm in B and C, and 50 μm in E. Arrows, Wolffian ducts; arrowheads, Müllerian ducts. (D) GFP intensity in the mesenchyme of the rostral and caudal reproductive tracts was quantified. (F) NIH3T3 cells were transfected with TCF/Lef-luc and CMV-β-gal plasmids and treated with 20 mM LiCl or conditioned media from SAM-HaCaT or shRNA-HaCaT cells and wild type or *Map3k1^ΔKD/ΔKD^* MEFs. Relative luc activities normalized by β-gal were calculated. Values are mean ± SEM. \*\**p*<0.01 and *** *p*< 0.001 are considered significantly different from the wild type embryo and the control media treatment.

At the most caudal end of the reproductive tract, the E-cadherin-positive and Pax2-negative UGS epithelium was GFP negative, but its surrounding mesenchyme exhibited strong GFP signals in E15.5 embryos of all three genotypes (Fig. 6A and 6E). In E13.5 embryos, both the UGS epithelium and mesenchyme are GFP negative in the wild type and the *Map3k1^ΔKD/ΔKD^* (Fig. S2G). Thus, the WNT activity in the UGS is independent of MAP3K1 activity.

To test whether MAP3K1 regulates secreted factors in the caudal MD epithelium to activate WNT in mesenchyme through a paracrine mechanism, we collected culture media from MAP3K1-compenent and deficient epithelial cells and Mouse Embryonic Fibroblasts (MEFs). The conditioned media were used to treat TCF/Lef-luc WNT reporter transfected NIH3T3 cells. Compared to un-treated cells, Lithium Chloride (LiCl), a known canonical WNT activator, produced a 2-fold increase in luciferase activity. Conditioned media derived from SAM also caused a significant increase of Luc activity by 1.5-fold, whereas media derived from shRNA cells did not have such an effect (Fig. 6E). Moreover, conditioned media of neither the wild type nor the *Map3k1^ΔKD/ΔKD^* MEFs were able to induce luc activity. Taken together, these data suggest that MAP3K1 induces secreted WNT activators in epithelial cells, not fibroblasts.

### MAP3K1 is required for optimal WNT7B expression

Several WNT ligands are known to be expressed in the internal genital epithelium and play pivotal roles in FRT development (Mullen and Behringer, 2014). We examined WNT ligand expression in E15.5 *Map3k1^+/ΔKD^*and *Map3k1^ΔKD/ΔKD^* embryos using RNAscope *in situ* hybridization. We detected, in a MAP3K1-independent manner, *Wnt7a* expression specifically in the MD epithelium and *Wnt5a* expression predominantly in the mesenchyme, whereas *Wnt4* expression occurred at low levels in both epithelium and mesenchyme of MD and WD (Fig. S3A-F). We also found that *Wnt7b* expression was unaffected by MAP3K1 ablation in the WD epithelium and the surrounding mesenchyme, but was significantly decreased in the *Map3k1^ΔKD/ΔKD^* MD epithelium (Fig. 7A and 7B). In cultured mouse keratinocytes, *Wnt7b* mRNA was reduced by 50% in *Map3k1^ΔKD/ΔKD^* compared to that in wild type cells (Fig. 7C). Likewise, the shRNA HaCaT cells displayed 40% reduction of *WNT7B* mRNA compared to the SAM cells (Fig. 7D). Thus, MAP3K1 deficiency down-regulates WNT7B expression in epithelial cells.

**Figure 7.**
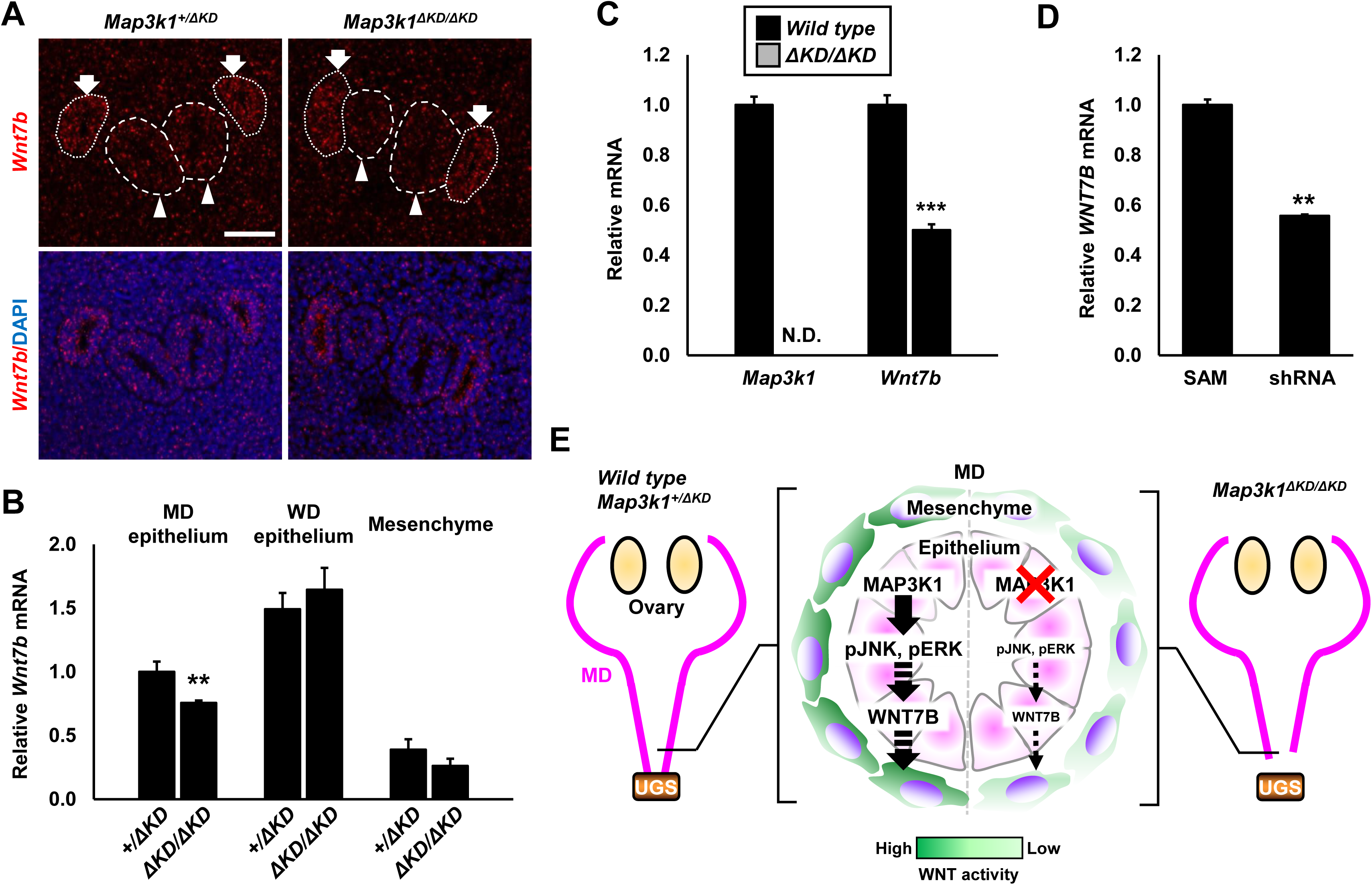
MAP3K1 deficiency downregulates *Wnt7b* expression in Müllerian duct (MD) epithelium. RNAscope *in situ* hybridization was performed using *Wnt7b* probe on transverse sections of the reproductive tract. (A) Representative images of *Wnt7b* in *Map3k1^+/ΔKD^* and *Map3k1^ΔKD/ΔKD^* female embryos at E15.5. Arrows, Wolffian ducts (WD); arrowheads, MD; scale bar, 50 μm. (B) Quantification of *Wnt7b* expression in MD and WD epithelium and mesenchyme of *Map3k1^+/ΔKD^* and *Map3k1^ΔKD/ΔKD^* female embryos at E15.5. (C) The expression level of *Map3k1* and *Wnt7b* mRNAs in wild type and *Map3k1^ΔKD/ΔKD^* keratinocytes. While the *Map3k1^ΔKD/ΔKD^* cells had no detectable *Map3k1* mRNA, they had significantly decreased expression of *Wnt7b*. (D) The level of *WNT7B* mRNA was lower in shRNA HaCaT cells than that in SAM cells. Values are mean ± SEM. \*\**p*<0.01 and \*\*\**p*<0.001 are considered significantly different from values in wild type, *Map3k1^+/ΔKD^* and SAM cells. N.D., not detected. (E) Working model for the role of MAP3K1 in female reproductive tract development. MAP3K1 induces JNK/ERK phosphorylation and Wnt7b expression in MD epithelium to maintain WNT signaling in the mesenchyme for caudal MD elongation; MAP3K1 deficiency reduces mesenchymal WNT activity, and causes defective MD caudal elongation and failure of MD-urogenital sinus (UGS) fusion.

## Discussion

Human genomics data have associated *MAP3K1* mutations with 46, XY DSDs, where the hyper-activated MAP3K1 perturbs sexual organ development in male patients (Ostrer, 2014; Pearlman et al., 2010). In this paper, we have used reverse genetics in the *Map3k1^ΔKD/ΔKD^* mice to explore the role of MAP3K1 in sexual development. We show that male mice carrying kinase-inactive MAP3K1 have normally formed reproductive organs and fertility, while female mice with the same kinase-inactive mutation exhibit defects in the caudal reproductive tract. The defects are traced to embryogenesis where the MDs in the *Map3k1^ΔKD/ΔKD^* embryos fail to extend caudally to connect with the UGS. This failure leads to distortion and complete absence of the MD-UGS junction in the neonatal homozygous knockout females, leading to imperforate vagina, reduced fertility and labor complications in the adult females. While all *Map3k1^ΔKD/ΔKD^* embryos have stunted MD, some of the neonates display morphologically normal female reproductive tracts, and 30% of the adult females are fertile. These observations suggest that the distorted embryonic MD may be amended at later developmental stages; the mechanism for this restoration is currently unknown. Together, the complementary mouse and human data strongly support a crucial role for MAP3K1 in sexual differentiation and sex organ development.

Analyses of MAP3K1-competent and -deficient epithelial cells derived from gene-editing of HaCaT cells and keratinocytes from wild type and *Map3k1^ΔKD/ΔKD^* mice indicate that WNT signaling is a downstream target of MAP3K1. Global gene expression, TCF/Lef-luc reporter analyses, and target gene expression studies in these cells suggest that MAP3K1 deficiency prevents optimal activation of WNT signaling. Correspondingly, compared to MAP3K1-competent cells, MAP3K1-deficient epithelial cells have reduced expression of *Wnt7b* and their conditioned media do not activate the WNT reporter in fibroblasts. *In vivo* data also support this view. *In vivo*, MAP3K1 is expressed in the epithelium of the reproductive tract, where it promotes the phosphorylation of JNK and ERK and the expression of *Wnt7b* in MD epithelium. MAP3K1 ablation impairs WNT signaling in the caudal MD mesenchyme. It is worth noting that the mesenchyme-specific β-catenin knockout mice also display stunted FRT (Deutscher and Hung-Chang Yao, 2007; MacDonald et al., 2009).

Wnt ligands play diverse roles in FRT differentiation and morphogenesis (Mericskay et al., 2004; Miller and Sassoon, 1998; Parr and McMahon, 1998; Prunskaite-Hyyrylainen et al., 2016; Vainio et al., 1999). Wnt7b is expressed in the FRT epithelium (Seishima et al., 2019) and we find that its expression is decreased in the MD epithelium of *Map3k1^ΔKD/ΔKD^*embryos. Wnt7b can act like Wnt4 to guide epithelial morphogenesis through a mechanism of epithelial-mesenchymal interaction (Kispert et al., 1998). Thus, reduced Wnt7b in *Map3k1^ΔKD/ΔKD^* embryos may contribute to WNT signaling deficiency, leading to MD regression similar to that displayed in WNT4-deficient mice and humans (Biason-Lauber et al., 2004; Prunskaite-Hyyrylainen et al., 2016).

The crosstalk of MAP3K1 and WNT is temporal-spatial in the embryonic reproductive tract. In the WNT reporter (TCF/Lef:H2B-GFP) embryos, MAP3K1 ablation does not change WNT activity in WD and MD epithelium, nor does it alter WNT activity in mesenchyme of the rostral reproductive tract and the UGS. Instead, MAP3K1 ablation reduces WNT activity specifically in mesenchyme of the caudal MD. Temporal-spatial WNT signaling activities along the rostral-caudal length regulate distinct aspects of reproductive tract differentiation and morphogenesis (Roly et al., 2018). The MAP3K1-dependent WNT activity in the MD mesenchyme likely contributes to the specific process of MD caudal elongation and MD-UGS fusion.

In summary, results presented here establish that MAP3K1 kinase activity is required for FRT development and that this function is mediated through MAP3K1-WNT crosstalk. Specifically, MAP3K1 regulates the expression of WNT ligands in the reproductive tract epithelium, which in turn activate WNT signaling in the surrounding mesenchyme that is essential for MD caudal elongation and MD-UGS fusion (Fig. 7E). Aberrant FRT development in humans leads to dysplasia or absence of the vagina, cervix and/or caudal uterus. These conditions affect approximately 5% of human females and are difficult to diagnose (Chan et al., 2011; Choussein et al., 2017). Clinical symptoms include, but are not limited to, failure to menstruate, periodic lower abdominal pain and a range of anomalies that affect multiple organs in the abdomen (Bombard and Mousa, 2014). Patients with this condition are at a greater risk of adverse reproductive outcomes, including reduced fertility, an increased miscarriage rate, preterm delivery, fetal malpresentation and complications at delivery (Chan et al., 2011). Our findings indicate that MAP3K1 loss-of-function is a molecular etiology contributing to congenital female reproductive defects in mice that is likely conserved across species.

## Materials and Methods

### Mice

Animals were housed in a pathogen-free vivarium in accordance with institutional policies. All animal experiments were approved in advance by the Institutional Animal Care and Use Committee at the University of Cincinnati College of Medicine. The *Map3k1^ΔKD^* mice were generated in house and described previously (Xia et al., 2000; Zhang et al., 2003). The Tg(TCF/Lef1-HIST1H2BB/EGFP)61Hadj (TCF/Lef:H2B-GFP) mice were from The Jackson Laboratory (JAX strain #013752)(Ferrer-Vaquer et al., 2010). The mice were backcrossed with C57BL/6 more than 12 generations to generate B6 congenic background. Tail genomic DNA was used for PCR genotyping.

### Cells, reagents and antibodies

HaCaT cells, a spontaneously immortalized human keratinocyte line, and NIH3T3 fibroblasts were obtained from ATCC. The Wild type and *Map3k1^ΔKD/ΔKD^* mouse keratinocytes and mouse embryonic fibroblasts (MEF) were prepared as described previously (Wang et al., 2021b; Zhang et al., 2003). The mouse keratinocytes were maintained in Defined Keratinocyte Serum Free Medium without calcium (K-SFM without calcium, Gibco) and routinely subcultured. The HaCaT, MEFs and NIH3T3 cells were cultured in Dulbecco’s Modified Eagle’s Medium (DMEM); the cell culture supplements added to DMEM, including fetal bovine serum (FBS), glutamine, nonessential amino acids, penicillin and streptomycin, were from Gibco. All other reagents and their sources are summarized in Supplemental Table 1; antibodies are listed in Supplemental Table 2.

### Modulating MAP3K1 expression in HaCaT cells

The *MAP3K1* shRNA lentiviral vectors were purchased from Sigma (Clone ID, TRCN000000616), and plasmids for CRISPR/Cas9 Synergistic Activation Mediator (SAM) system, i.e. lenti-sgRNA(MS2)_puro, dCas9VP64, and MS2-P65-HSF1 were purchased from Addgene (Addgene, cat. #73797, 61425, and 61426). The single guide RNAs (sgRNA) for *MAP3K1* were designed based on publicly available filtering tools (https://zlab.bio/guide-design-resources) (Suppl. Table 3); the sgRNA was cloned into lenti-sgRNA(MS2)_puro. The lentiviral vectors were co-transfected with packaging psPAX2 (Addgene #12260) and envelope pMD2.G (Addgene #12259) plasmids into 293T packaging cells (ATCC). Viral production, amplification and titration were done following the protocol in Addgene.

To make MAP3K1 deficient HaCaT, the cells were transduced with lentivirus containing *MAP3K1* shRNA and then selected with puromycin to obtain stable “shRNA” cells. To make MAP3K1 over-expression HaCaT, the SAM system was used (Konermann et al., 2015). Specifically, the lentiviruses for sgRNA, dCas9VP64 and MS2-P65-HSF1 were used to transduce the HaCaT cells, which were subsequently selected with 3mg/ml puromycin, 10 mg/ml blasticidin and 5 mU hygromycin in DMEM containing 5% FBS to generate the stable “SAM” cells.

### Histology, Immunohistochemistry and RNAscope *in situ* hybridization

The lower body of appropriately euthanized embryos and neonates were collected and fixed by immersion in 4% paraformaldehyde (PFA, Thermo Fisher Scientific) for overnight at 4 °C. For microscopic analyses, the fixed tissues were embedded routinely in paraffin, sectioned serially at 5 µm, and conventional hematoxylin and eosin (H&E) staining. Images were captured by bright-field light microscopy with different magnifications.

For immunohistochemistry, fixed tissues were immersed with sucrose gradients (15% and 30%), covered in Tissue-Tek O.C.T. Compound, flash-frozen on dry ice, and stored at -80 ℃. Serial 12-μm-thick sections were obtained and washed in PBS, immersed in boiled 10 mM citrate buffer (pH 6.0) for 30 min, blocked with PBS containing 5% bovine serum albumin (BSA) for 1 h at R.T, and incubated with a diluted primary antibody (Supplemental Table 2) overnight at 4℃ followed by secondary antibodies and Hoechst 33342 staining.

RNAscope *in situ* hybridization was done as described previously (Wang et al., 2012) and following the manufacturer’s instructions. Briefly, 12-µm-thick sections were deparaffinized and washed. After epitope retrieval in boiled citrate buffer (10 mM, pH 6.0) for 30 min, the sections were incubated with the commercial synthetic probes (Supplemental Table 1) at 40℃, followed by signal amplification and fluorescent dye labeling. Images were captured by the Zeiss Axio fluorescent microscope.

Microscopic evaluation of the adult FRT was done at the Comparative Pathology & Mouse Phenotyping Shared Resource of The Ohio State University College of Veterinary Medicine. At the necropsy of 1- to 3-month-old female mice, animals were appropriately euthanized, and the reproductive tract and selected major viscera were removed and fixed by immersion in neutral buffered 10% formalin. The samples were processed routinely into paraffin, sectioned at 5 µm, stained with H&E, and evaluated by ACVP board-certified veterinary pathologists.

### Whole mount tissue immunofluorescence staining and X-gal staining

A previously described protocol was adapted for whole-organ immunofluorescence (Messal et al., 2021). Briefly, the lower bodies of embryos and neonates were fixed with 4% PFA overnight at 4 °C. For antigen retrieval, the tissues were incubated with FLASH reagent 2 (200 mM borate buffer, 250 g/liter urea and 80 g/liter Zwittergent) for 1 h at RT, followed with overnight incubation in FLASH reagent 2 at 55℃. For immunostaining, the tissues were washed with PBST (PBS plus 0.2% Triton X-100) for 30 min at RT, immersed in blocking buffer {PBST with 10% (wt/vol) FBS, 0.02% (wt/vol) sodium azide, 1% (wt/vol) BSA and 5% (vol/vol) DMSO} for 1 h at RT, and subjected to immunolabelling with primary and secondary antibodies (Supplemental Table 2) for 5 and 3 days, respectively, at RT. For tissue clearing, after extensive washing, tissues were dehydrated gradually in 30%, 50%, 75% and 100% (vol/vol) methanol, followed by a serial gradient with 25% - 100% of Benzyl Alcohol/ Benzyl Benzoate (BABB). Cleared tissues were placed in chamber slides, and imaged using a Zeiss confocal microscope (LSM700).

For whole mount X-gal staining, the lower bodies of wild type and *Map3k1^+/ΔKD^* embryos were collected and fixed with a solution containing 2% PFA, 0.2% glutaraldehyde, 5 mM EGTA, 2 mM MgCl_2_ an 0.02% NP40 for 20 min at RT. The tissues were immersed in staining solution, containing 100 mM phosphate buffer, 5 mM K_3_Fe(CN)_6_, 5 mM K_4_Fe(CN)_6_, 2 mM MgCl_2_, 0.02% NP40 and 1 mg/mL X-gal overnight at 37℃. The tissues were post-fixed with 4% PFA at 4℃ overnight, processed through sucrose gradient, embedded in OCT flash-frozen, and sectioned at 12 µm. The sections were counter stained with diluted hematoxylin for 5 min at R.T. After dehydration through ethanol and xylene, sections were covered in Mounting Medium Xylene, and images were captured under a bright-field light microscopy with different lenses.

### Proliferation and apoptosis detection

Embryos were harvested from pregnant mice given an intraperitoneal injection of EdU (5 mg/kg b.w.) 2 hours prior to collection. The embryos were fixed, embedded and sectioned as described above for immunohistochemistry. Cell proliferation was determined with the iClick EdU Imaging Kit (ABP Biosciences) and apoptosis was measured using the ApopTag Peroxidase *in Situ* Apoptosis Detection Kit (Millipore Sigma), following the manufacturers’ protocols. Images were captured with a Zeiss Axio microscope. The positively stained cells were quantified from at least 3 embryos/genotype in 4 sections per embryo.

### Transfection and reporter assays

Transfection of the NIH3T3, HaCaT SAM and shRNA cells were done using Lipofectamine 2000 (Invitrogen). Briefly, cells were seeded in 24-well tissue culture plates and transfected with TRE-luc and TCF/Lef-luc, together with the ß-galactosidase plasmids following the manufacturer’s instructions.

Serum-free DMEM incubated for 24 h with shRNA and SAM HaCaT cells, and wild type and *Map3k1^ΔKD/ΔKD^* MEF was collected as “conditioned media”. The *in vitro* studies of epithelial-mesenchymal interaction were performed by measuring the TCF/Lef-luc activity in transfected NIH3T3 cells. The transfected cells were treated with the conditioned media for 24 h, and cell lysates were examined for luciferase activities with the luciferase assay system kit (Promega). The relative luciferase activities were calculated after normalization with the ß-gal activities. Results represent duplicate data from at least 3 biological samples of each genotype.

### RNA isolation, reverse transcription and real-time RT-PCR

Total RNA was extracted from cultured cells and purified with PureLink® RNA Mini kit (Invitrogen). The RNA (0.5 μg) was subjected to reverse-transcription using the SuperScript™ IV Reverse Transcriptase kit (Invitrogen). The cDNA was subjected to quantitative real time PCR (qRT-PCR) using MX3000p thermal cycler system and SYBR Green QPCR Master Mix. Relative gene expression was calculated by the comparative ΔΔCt-method normalized to a constitutively expressed housekeeping gene (*GADPH*). All determinations were performed with triplicate samples. Primer sequences are listed in Supplemental Table 3.

### RNA-seq and pathway analyses

Total RNA was isolated from SAM and shRNA HaCaT cells and wild type and *Map3k1^ΔKD/ΔKD^* mouse keratinocytes. The RNA quality was QC analyzed by Bioanalyzer (Agilent, Santa Clara, CA). RNA sequencing was performed with biological triplicate samples by the Genomics, Epigenomics and Sequencing Core at the University of Cincinnati using established protocols described previously (Qiu et al., 2023; Wang et al., 2021b), and in the Supplemental Materials and Methods. The RNA-seq data were deposited at Gene Expression Omnibus (GEO) publicly accessible (GSE39240). The differentially expressed genes were subjected to Bioinformatics analyses using Ingenuity Pathway Analyses software (IPA, Qiagen).

### Image quantification and statistical analyses

The ratio of EdU- or TUNEL-positive cells versus the total epithelial cells in MD and WD was calculated. The immunofluorescent intensity (i.e., immunohistochemistry, GFP and RNAscope) was measured in MD and WD epithelium and surrounding mesenchyme. The WD and MD areas consisting of epithelial cells expressing Pax2 or E-cadherin were outlined and the mean intensity values of target proteins/genes were measured. Quantification was done by Image J software (ver. Fiji-2.1.1). The level of the targets in the images was determined after background subtraction. At least three embryos/genotype were used for quantification. All data are shown as mean ± SEM. Statistical comparisons were performed with Student’s t-test or one-way analysis of variance following Tukey’s multiple comparison test. Values of *p* < 0.05 (*), *p* < 0.01 (**), and *p* < 0.001 (***) were considered to be statistically significant.

## Acknowledgements

We thank Dr. Yueh-Chiang Hu at the Cincinnati Children’s Hospital Medical Center, and Drs. Divaker Choubey and Chia-I Ko at the University of Cincinnati for reagents and technical assistance. We also thank Dr. Bing Su at Shanghai Jiaotong University, who brought our attention to the IPV phenotype in *Map3k1_ΔKD_* mice. Work is supported in part by NIH grants RO1HD098106, R21ES033342, RO1ES010807 and P30ES006096.

